# Identification and phenotype of MAIT cells in cattle and their response to bacterial infections

**DOI:** 10.1101/2020.11.09.374678

**Authors:** Matthew D. Edmans, Timothy K. Connelley, Siddharth Jayaraman, Christina Vrettou, Martin Vordermeier, Jeffrey Y. W. Mak, Ligong Liu, David P. Fairlie, Emmanuel Atangana Maze, Tiphany Chrun, Paul Klenerman, Sidonia B. G. Eckle, Elma Tchilian, Lindert Benedictus

**Author notes:** **Correspondence:** Matthew Edmans. These authors have contributed equally to this work (Paul Klenerman, Sidonia B. G. Eckle, Elma Tchilian, Lindert Benedictus).

## Abstract

Mucosal-associated invariant T (MAIT) cells are a population of innate-like T cells that utilise a semi-invariant T cell receptor (TCR) α chain and are restricted by the highly conserved antigen presenting molecule MR1. MR1 presents microbial riboflavin biosynthesis derived metabolites produced by bacteria and fungi. Consistent with their ability to sense ligands derived from bacterial sources, MAIT cells have been associated with the immune response to a variety of bacterial infections, such as *Mycobacterium spp*., *Salmonella spp. and Escherichia coli*. To date, MAIT cells have been studied in humans, non-human primates and mice. However, they have only been putatively identified in cattle by PCR based methods; no phenotypic or functional analyses have been performed. Here, we identified a MAIT cell population in cattle utilising MR1 tetramers and high-throughput TCR sequencing. Phenotypic analysis of cattle MAIT cells revealed features highly analogous to those of MAIT cells in humans and mice, including expression of an orthologous TRAV1-TRAJ33 TCR α chain, an effector memory phenotype irrespective of tissue localisation, and expression of the transcription factors PLZF and EOMES. We determined the frequency of MAIT cells in peripheral blood and multiple tissues, finding that cattle MAIT cells are enriched in mucosal tissues as well as in the mesenteric lymph node. Cattle MAIT cells were responsive to stimulation by 5-OP-RU and riboflavin biosynthesis competent bacteria *in vitro*. Furthermore, MAIT cells in milk increased in frequency in cows with mastitis. Following challenge with virulent *Mycobacterium bovis*, a causative agent of bovine tuberculosis and a zoonosis, peripheral blood MAIT cells expressed higher levels of perforin. Thus MAIT cells are implicated in the immune response to two major bacterial infections in cattle. These data suggest that MAIT cells are functionally highly conserved and that cattle are an excellent large animal model to study the role of MAIT cells in important zoonotic infections.

## 1 Introduction

Mucosal-associated invariant T (MAIT) cells represent the largest antigen specific α/β T cell population in humans, comprising up to 10% of all T cells in the periphery and 50% in the liver (1-3). Unlike conventional α/β T cells that recognize peptides presented by MHC class I (MHC-I) or MHC class II (MHC-II) molecules, MAIT cells recognise microbial riboflavin biosynthesis derived metabolites presented by a monomorphic MHC-I like molecule, MHC related protein 1 (MR1) (4-6). To date the most potent ligand presented by MR1 is 5-(2-oxopropylideneamino)-6-D-ribitylaminouracil (5-OP-RU), which is a derivative of a key riboflavin biosynthetic intermediate (7, 8). MAIT cells are unconventional T cells that emerge from the thymus in a “pre-primed” state and express an effector memory phenotype (in humans CD45RA^−^CD45RO^+^CD95^hi^CD62L^lo^)(9). This facilitates rapid innate-like responses to bacterial infections, such as *Francisella tularensis* (10), *Legionella longbeachae* (11) and *Mycobacterium tuberculosis* (TB) (12, 13).

MR1 is estimated to have appeared around 170 Mya ago in a common ancestor of marsupial and placental mammals (14, 15) and is the most highly conserved MHC molecule in its ligand-binding α1-/α2-domains (14, 16-18). Despite this, MR1 has been lost in a select number of species, including carnivores and lagomorphs (15). MAIT cells express an essentially invariant T cell receptor (TCR) α chain (TRA), composed in humans of the variable gene segment 1-2 (*TRAV1-2*) (also known as Vα7.2 in Arden nomenclature) rearranged with the TRA joining gene segment 33 (*TRAJ33*) (also known as Jα33), or less commonly *TRAJ12* (Jα12) and *TRAJ20* (Jα20), paired with TCR β chains (TRB) featuring dominant usage of specific TRBV gene segments (12, 19-22). Species that possess the MR1 gene also express a homologue of the canonical MAIT TCR *TRAV1-2* gene segment (15) and the entire canonical MAIT TCR α chain is conserved across mammalian species, with homologous TCR α chains detected in mouse (20), cattle (20, 23), sheep (23), pigs (24) and macaques (25). MAIT cells follow similar developmental pathways in humans and mice (26, 27) however, mice have much smaller MAIT cell populations than humans with differences in phenotype and tissue distribution (26). Thus, whilst mice have provided valuable information on MAIT cells in protection against infections (1, 11, 28, 29), pathology (30) and tissue repair (31), this might not always be fully representative of MAIT cell function in humans.

Beyond primates and mice, information on MAIT cells is limited. Cattle are an economically important livestock species and are also a relevant large animals model for human infections, including tuberculosis (32, 33) and respiratory syncytial virus (RSV) (34). Cattle express the MR1 gene and the canonical MAIT cell TRA (17, 20, 23, 35). However, MAIT cells have not been characterised directly and there is no knowledge of the phenotype and function of MAIT cells in cattle. In humans and mice, as well as more recently in macaques, fluorescently labelled MR1 tetramers loaded with the MAIT cell activating ligand 5-OP-RU have become the gold standard to identify MAIT cells (7, 19, 36-40). Tetramers loaded with the MR1 ligand 6-formylpterin (6-FP) (6, 41) or its acetylated analogue, acetyl-6-FP (Ac-6-FP) (41), typically do not bind to MAIT cells and are often used as negative controls for MR1-5-OP-RU tetramer staining in humans (42). Here we used human MR1 tetramers and synthetic 5-OP-RU antigen to identify and characterise MAIT cells in cattle. Further, we show that cattle MAIT cells can be activated by bacteria *in vitro* and that MAIT cells respond in the context of mastitis and *Mycobacterium bovis* (*M. bovis*) infection in cattle, suggestive of a role for MAIT cells in these diseases, caused by riboflavin biosynthesis competent pathogens.

## 2 Material and Methods

### 2.1 Animals

All animal experiments were conducted within the limits of a United Kingdom Home office license under the Animal (Scientific Procedures Act 1986) (ASPA) and were reviewed and approved by the Animal Welfare and Ethical Review Bodies of the institutes where the experiments were performed (the Roslin Institute and the Animal and Plant Health Agency). Sampling milk from cattle is below the threshold of pain, suffering, distress or lasting harm that requires A(SP)A licensing and the procedure was reviewed by the Veterinary Ethical Review Committee of the Royal Dick School of Veterinary Studies (RDSVS), Edinburgh University.

Healthy, Holstein-Frisian cattle aged between three to fifty-six months were housed at the Edinburgh University farms or at the Animal Plant and Health Agency (APHA) facilities at Weybridge. Blood was sampled from the jugular vein and peripheral blood mononuclear cells (PBMC) were isolated from blood by density gradient centrifugation and cryopreserved. To harvest tissues, seven male cows (aged 10, 10, 10, 10, 20, 55 and 56 months) were culled by schedule 1 methods under the ASPA followed by auscultation of the heart to confirm cessation of the circulation.

The BCG-vaccination and *M. bovis*-challenge study was described previously (43). In short, six bovine tuberculosis-free 6-months old male Holstein-Friesian (cross) calves were vaccinated subcutaneously with 4.6×10^6^ CFU *M. bovis* BCG Danish SSI 1331 (Statens serum Institute) and two calves served as controls. Nine weeks later all calves were infected with 10^4^ CFU virulent *M. bovis* AF2122/97 via the endobronchial route. Twenty weeks post BCG vaccination all animals were euthanised and post-mortem examination was performed as described by Vordermeier et al. (44). Gross visible lesions of lungs and the lymph-nodes of the head and pulmonary regions were scored semi-quantitatively resulting in a total gross pathology score. Blood was sampled regularly and PBMC were isolated and cryopreserved.

Milk was sampled from Holstein-Friesian cattle housed at Langhill Dairy Farm, the teaching farm of the RDSVS, Edinburgh University.

### 2.2 Tissue sampling and processing

Single cell suspensions were obtained from peripheral blood, prescapular lymph node (Ln), mesenteric Ln, lung, bronchial alveolar lavage (BAL), ilium, spleen, liver, and milk. Peripheral blood was diluted 1:1 in PBS and layered over Histopaque-1077 (Sigma-Aldrich) before centrifugation at 1200 g for 20 minutes. Cells were washed and resuspended in RPMI supplemented with 10% FCS and 1% penicillin streptomycin (Sigma Aldrich) (complete media) or in PBS buffer supplemented with 2% FCS and 0.01% Azide. Prescapular Ln and mesenteric Ln were suspended in complete media before being manually disrupted and passed through a 100 μM cell strainer. For lung and BAL, a lung lobe was removed and the main bronchus washed with 750 ml of PBS. Lungs were massaged for 30 seconds before BAL fluid was collected. The BAL fluid was transferred into 50 ml Falcon tubes and centrifuged at 400 g for 10 minutes and resuspended in complete media. A piece of lung was dissected in ∼0.5 cm cubes and resuspended in 7 ml serum free RPMI containing 30 μg/ml DNAse and 700 μg/ml collagenase (Sigma Aldrich) and in C Tubes (Miltenyi Biotec) disassociated using the gentleMACS Octo Dissociator (Miltenyi Biotec). The C tubes were then incubated for 60 minutes at 37 °C, before being dissociated a second time, cells re-suspended in complete media and passed through a 100 μM cell strainer. Ileum, spleen and liver were also dissected and, samples were suspended in C Tubes in complete media and disassociated using the gentleMACS Octo Dissociator. Following disassociation, the resulting ileum and spleen cell suspension was passed through a 100 μM cell strainer. For liver, the cell suspension was resuspended in a 50 ml Falcon tube in 20 ml 35% PERCOL which had previously been made isotonic with 10x PBS and diluted with complete media. The 35% PERCOL was underlayed with 10 ml 70% PERCOL and centrifuged at 1200 g for 20 minutes. Cells were collected at the interface and resuspended. For all tissues other than liver and peripheral blood, the obtained cell suspensions were layered onto Histopaque-1077 (Sigma-Aldrich) before centrifugation at 1200 g for 20 minutes and the lymphocytes collected at the interphase. All cells were finally filtered through a 100 μM cell strainer, red blood cells lysed with an Ammonium Chloride Lysis buffer if required, washed and if not immediately used for assays, cryopreserved in FCS containing 10% DMSO. To isolate cells from milk, milk was centrifuged (400 g, 4 °C, 15 minutes). The resulting fay layer was removed with a pipette tip and the supernatant discarded. The pellet was resuspended in PBS and moved to a clean tube and the procedure was repeated. For the second PBS wash, the cell suspension was filtered with a 70 μM strainer and after centrifugation the cell pellet was resuspended in PBS + 2% FCS for downstream procedures.

### 2.3 ELISPOT

Frequencies of IFN-γ secreting cells were determined by ELISPOT IFN-γ assay. MultiScreen-HA ELISPOT plates (Merck Millipore) were coated with primary anti-IFN-γ clone CC330 (Serotec, 2 μg/ml) and incubated at 4°C overnight. Plates were washed and blocked with complete media for two hours. Plates were seeded with 2.5 × 10^5^ PBMC and stimulated with either 1 μM 5-OP-RU (produced in house as previously described (8)), 4 μg/ml ConA (Sigma-Aldrich) or medium control. Plates were incubated overnight at 37 °C before washing with PBS containing 0.05% Tween 20 and addition of secondary biotinylated IFN-γ detection Ab (clone CC302 (Serotec, 2 μg/ml)). Plates were incubated for 2 hours at room temperature, washed a further five times, and streptavidin–alkaline phosphatase (Invitrogen) was added for 1 hour. Spots were visualised using alkaline phosphatase substrate kit (Bio-Rad) and the reaction stopped using water. Immunospots were enumerated using the AID ELISPOT reader (AID Autoimmun Diagnostika). Results are expressed as the total number of IFN-γ producing cells per 10^6^ input PBMC following subtraction of the average number of IFN-γ positive cells in medium control wells.

### 2.4 MR1 tetramers

The MR1 tetramer technology was developed jointly by Dr. James McCluskey, Dr. Jamie Rossjohn, and Dr. David Fairlie (7) and the human MR1 tetramers (human MR1-5-OP-RU and human MR1-6-FP) were obtained from the NIH Tetramer Core Facility as permitted to be distributed by the University of Melbourne.

### 2.5 Flow cytometry

For phenotyping, *ex vivo* isolated or thawed cryopreserved cells were seeded into a 96 well plate at 1-3 × 10^6^ cells / well. Cells were stained with pre-diluted tetramer in PBS + 2% FCS for 40 minutes at room temperature. Following tetramer staining, primary antibodies (Table 1) were added in PBS buffer supplemented with 2% FCS and 0.01% Azide, and Near-Infrared or Yellow Fixable LIVE/DEAD stain (Invitrogen or Molecular probes) for 15-30 minutes at 4 °C. If required, cells were washed, and secondary antibodies added for 15-30 minutes at 4 °C. Cells were resuspended in buffer supplemented with 2% FCS and 0.01% sodium Azide and either immediately analysed or fixed in 4% paraformaldehyde and resuspended in PBS prior to analysis.

**Table 1.**
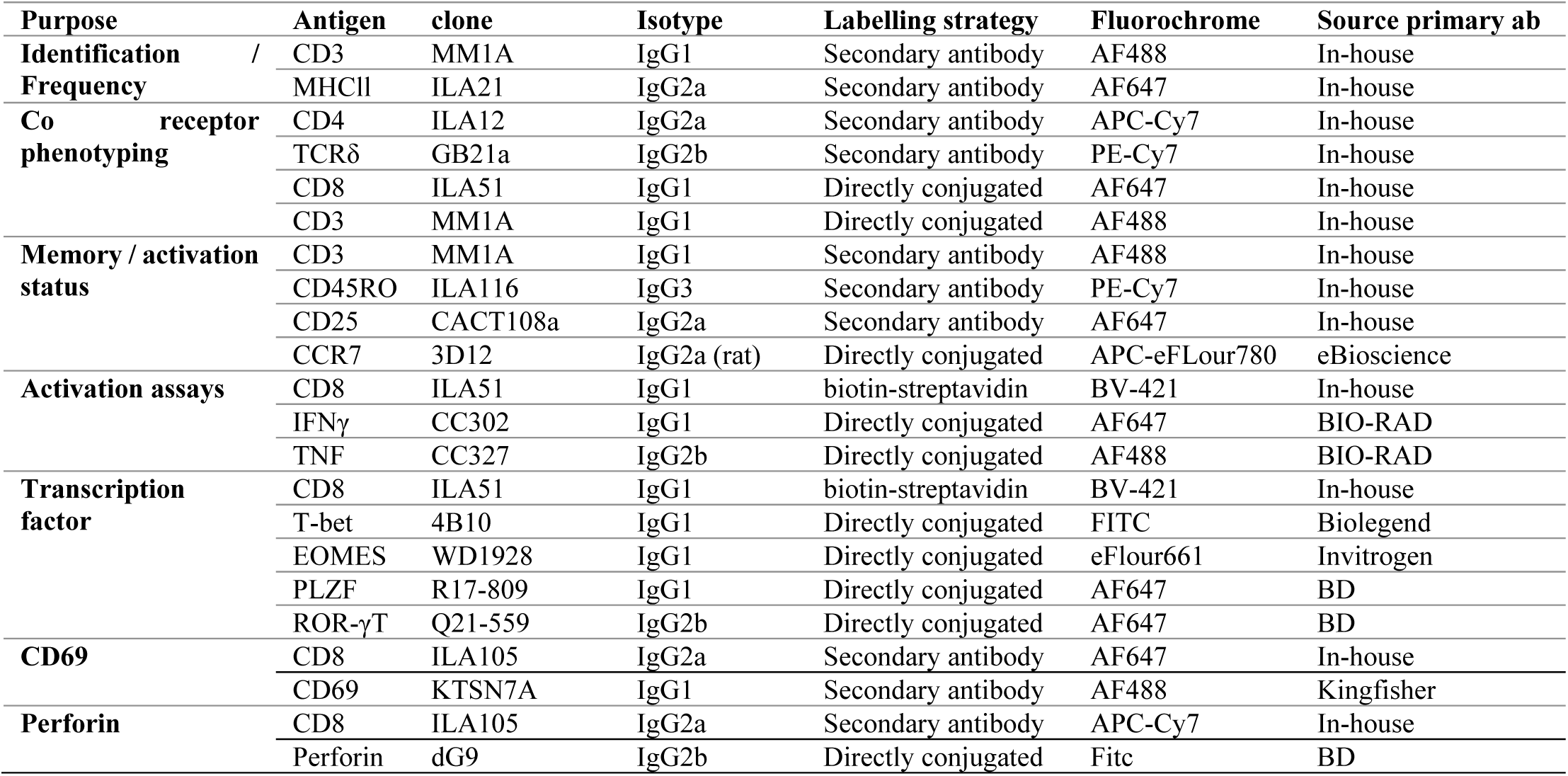
Monoclonal antibodies used in flow cytometry experiments.

For activation experiments, cryopreserved PBMC were thawed and stimulated with titrating amounts of 5-OP-RU, fixed *E*.*coli* or *S. aureus* at 30 bacteria per cell (BpC) for 7 hours. Stimulation with human IL-1212 (Miltenyi) and IL-18 (Biolegend), either alone (50 ng/ml) or in combination (50 ng/ml), and *M. bovis* strain BCG (3BpC) were performed over 18 hours. For mitogen stimulation, PMA and Ionomycin (eBioscience cell stimulation cocktail) were added to cells for 6 hours. Ac-6-FP (Schircks Laboratories) was used as a blocking reagent for some 5-OP-RU stimulations and was added 30 minutes prior to 5-OP-RU. For all stimulation conditions, Golgi plug (BD Biosciences) was added for the final 6 hours of stimulation. Cells were stained with human MR1-5-OP-RU tetramer for 40 minutes at RT in the dark prior to the addition of anti-bovine CD8-biotin (Clone ILA51) and Near-Infrared Fixable LIVE/DEAD stain (Invitrogen or Molecular Probes) for 20 minutes at 4 °C. BV421 conjugated streptavidin was added for 20 minutes at 4 °C prior to fixation and permeabilisation (BD Fix perm kit). Anti IFN-γ-FITC (clone CC302) and TNF-APC (clone CC327) were added for 20 minutes at 4 °C. Cells were washed and re-suspended in PBS prior to analysis.

For transcription factor stains, thawed PBMC were stained with tetramer, live/dead stain and primary and secondary antibodies as described. Cells were fixed in Fix perm buffer (eBioscience) for one hour followed by 1 hour in perm/wash buffer (BD). Conjugated antibodies specific for transcription factors PLZF (clone R17-809), EOMES (clone WD1928), ROR-γT (clone Q21-559) and T-bet (clone 4B10) diluted in perm/wash buffer were added for 1 hour at 4 °C. Cells were resuspended in PBS prior to analysis. Positive staining for each marker was determined by appropriate isotype controls and non-MAIT cell control populations.

All cells were acquired on a MACSquant analyser 10 (Miltenyi) or LSRFortessa (BD Biosciences).

### 2.6 Bovine T cell receptor α and β chain sequencing

To obtain CD8^+^MR1-5-OP-RU tetramer^-^ (non-MAIT) and CD8^+^MR1-5-OP-RU tetramer^+^ (MAIT) cell populations for TCR sequencing, sequential MACS isolation and cell sorting were performed. Freshly isolated PBMC from four 13 months old heifer calves were stained with biotinylated mAb ILA51 (anti-CD8, produced in-house) in PBS supplemented with 0.5% FCS and 2 mM EDTA for 15 minutes on ice with resuspending every 5 minutes. Following washing, cells were stained with magnetic anti-biotin beads (Miltenyi Biotec) and labelled cells were isolated using MS columns (Miltenyi Biotec) according to the manufacturer’s instruction. Isolated cells were stained with MR1-5-OP-RU tetramer, followed by staining with mAbs MM1a (anti-CD3) and ILA105 (anti-CD8) and secondary antibodies and Fixable Yellow Dead Cell Stain (Thermofisher Scientific). These cells were sorted using a FACSAria III (BD), gating on FSC and SSC, singlets and live cells, respectively. Within these gates CD8^+^MR1-5-OP-RU^-^ and CD8^+^MR1-5-OP-RU^+^ T cell populations were sorted directly into lysis buffer for RNA isolation (RNeasy Plus Micro kit, Qiagen). Purity of ungated sorted CD8^+^humMR1-5-OP-RU^+^ cells was between 91-99% (Fig. S1). RNA was isolated from 20,000 cells using the RNeasy Plus Micro kit according to manufacturer’s instruction, with addition of 4 ng/μl carrier RNA. cDNA was generated using the SuperScript IV kit (ThermoFisher Scientific) with a SMART oligo containing unique molecular identifiers (AAG CAG UGG TAU CAA CGC AGA GTUNNNNUNNNNUNNNNUCTTggggg (where N represents a mixture of A, T, G and C and lower case ‘g’ represents RNA bases) and the uracil-containing primers subsequently removed by treatment with UDG (NEB, Hitchin, UK). TRA and TRB sequences were amplified using a pair of 5’ ‘step-out’ primers specific for the SMART oligo (long 5’ primer - CTA ATA CGA CTC ACT ATA GGG CAA GCA GTG GTA TCA ACG CAG AGT, and short 5’ primer - CTA ATA CGA CTC ACT ATA GGG CAA GCA G) in combination with *TRAC*-(TGG GGT TGG GGT CCT TGA CTT) and *TRBC*-(GAC SYG GCT CAG ATC ATC) specific 3’primers. PCR amplification was conducted using Phusion HF reagents (NEB) and employed the following cycling conditions: manual hot start, 30 seconds at 98 °C, 35 cycles of (98 °C for 10s, 65 °C for 30s, 72 °C for 30s) and a final extension period of 5 minutes at 72 °C. PCR products of the anticipated size were excised following agarose gel electrophoresis, purified using AMpure beads (Beckman Coulter, Indiana, US) and sent to Edinburgh Genomics for sequencing on the Illumina MiSeq v3. Platform. Analysis of the TCR repertoire was conducted with the MiXCR package (45) using an external database of bovine TRAV, TRBV, TRAJ and TRBJ genes derived from a combination of published (46, 47) and unpublished data.

### 2.7 Bacteria for MAIT cell stimulation

Tryptic soy broth (TSB) was inoculated with *Escherichia coli* strain DH5α and *Staphylococcus aureus* strain RF122 from glycerol stocks and cultured overnight at 37 °C and 200 RPM. The next morning a 1:100 dilution in TSB was cultured for two hours and OD at 600 nm measured to estimate CFU/ml (OD 600 nm of 1.0 = 8 × 10^8^ CFU/ml). Bacteria were washed with PBS, fixed in 1% formaldehyde for 5 minutes at RT, washed thrice with PBS, resuspended in CM and stored at -20 °C.

*M. bovis* strain BCG Danish SSI 1331 (Statens Serum Institute) was cultured in glass vials in Middlebrook 7H9 broth supplemented with Tween 80, Amphotericin B (all Sigma-Aldrich), and BD Difco™ BBL™ Middlebrook ADC Enrichment and incubated at 37 °C with agitation with a magnetic stir bar. After 25 days the bacterial culture was vortexed vigorously and after 1 minutes the ‘supernatant’ was harvested and passed several times through a 21g needle to obtain a single bacteria suspension. Bacteria were pelleted and resuspended in 7H9 broth supplemented with 30% glycerol, aliquoted and stored at -80 °C. Thawed aliquots were serially diluted 1:10 in 7H9 broth and 100 μl of these suspensions were cultured on Middlebrook 7H11 agar, supplemented with OADC (both BD), to determine CFU/ml.

### 2.8 Data analyses and statistics

Flow cytometry data were analysed using FlowJo v10. Descriptive and statistical analyses were performed using Prism software version 8 (GraphPad). Data presented in the text and figures are means with standard error of the mean (SEM). P-values corrected for multiple comparisons ≤0.05 were considered significant. * p≤0.05, ** p<0.01, *** p<0.001, **** p<0.0001.

## 3 Results

### 3.1 Identification of MAIT cells in cattle

Due to the high level of MR1 (14, 15, 17, 35) and MAIT TCR α chain (14, 15, 20, 23, 24, 40) conservation between species, we hypothesised that human MR1 tetramers would likely cross react with cattle MAIT cells. We isolated PBMC from a cohort of cattle (n = 17) of varying age (3 to 56 months) and indeed staining with MR1-5-OP-RU tetramers identified a clear population of CD3^+^ tetramer^+^ ‘putative’ MAIT cells (**Fig. 1A-B**) with a mean frequency of 0.6% amongst CD3^+^ cells, which was comparable to previous qPCR estimates of MAIT cell frequency in cattle (∼0.2% of transcribed TRA) (23). A much lower frequency was identified by the control MR1-6-FP tetramer (**Fig. 1A-B**). The frequency of MAIT cells varied greatly between individuals with a range of 0.18-1.72% and an interquartile range (IQR) of 0.33-0.66% of total T cells. Within this age cohort (3-56 months), there was no correlation between age and MAIT cell frequency. Whilst in humans MAIT cells make up a higher proportion of T cells (mean 3.1%), frequencies of MAIT cells in humans also vary widely between individuals with an IQR of 1.3-4.5% (38). The most potent MAIT cell ligand identified to date is 5-OP-RU (7, 8), which specifically induces cytokine secretion, including IFN-γ, in MAIT cells but not in other T cells (7, 41). To corroborate the identification of MAIT cells using tetramers, we next determined whether a 5-OP-RU reactive population was present in cattle PBMC by IFN-γ ELISpot following stimulation with synthetic 5-OP-RU (8) (**Fig. 1C**). Following stimulation with 5-OP-RU a mean of 125 IFN-γ secreting cells /10^6^ PBMC were detected, demonstrating that there was a 5-OP-RU reactive T cell population in cattle. In summary, we identified a population of MR1-5-OP-RU tetramer^+^ T cells and 5-OP-RU reactive cells in cattle peripheral blood, strongly suggesting that we identified a MAIT cell population in cattle.

**Figure 1:**
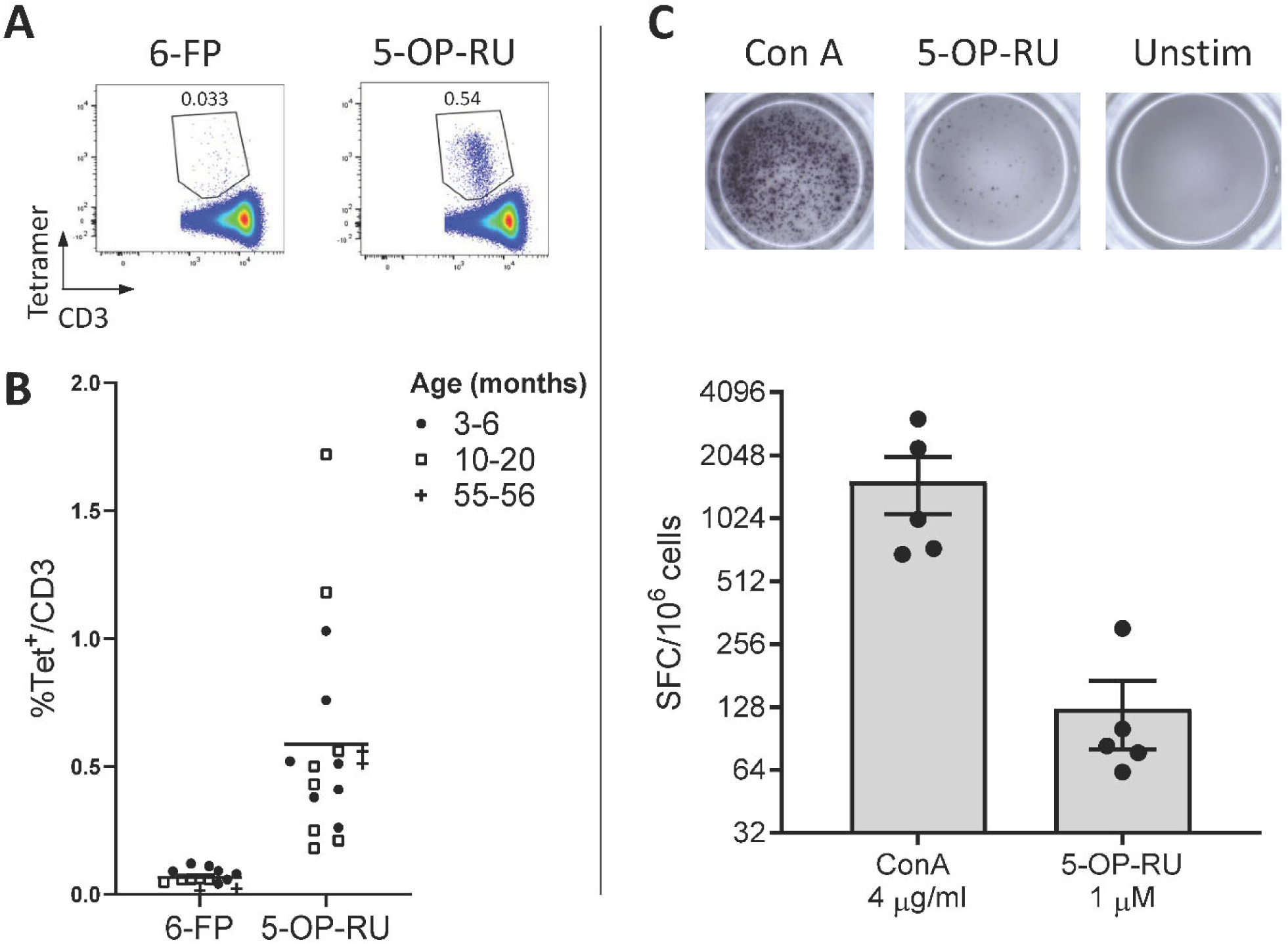
Identification of MAIT cells in cattle. **(A)** Representative flow cytometry plots depicting human MR1-6-FP and MR1-5-OP-RU tetramer staining of bovine PBMC, gated on CD3^+^ cells. **(B)** Fraction of human MR1-5-OP-RU (mean 0.59, range 0.18-1.72, n = 17) and MR1-6-FP (mean 0.06, range 0.01-0.12, n = 14) tetramer^+^ cells within MHC-II^-^ CD3^+^ PBMCS from cows of various ages. **(C)** Representative images and summarized data (mean ± SEM, n = 5) of IFN-γ secreting cells following stimulation of PBMC with 5-OP-RU and Concanavalin A (ConA) as a control. Responses are medium control subtracted.

### 3.2 Phenotype of MAIT cells in bovine peripheral blood

MAIT cells are unconventional T lymphocytes with functional and phenotypic features that distinguishes them from conventional T lymphocytes, including an effector memory phenotype prior to antigen exposure (48), enrichment in mucosa (2) and expression of specific transcription factors such as PLZF (49). According to co-receptor expression (**Fig. 2A**), cattle peripheral blood MAIT cells were predominantly CD8^+^ (mean 73.9%, IQR 64-87%) or double negative (mean 19.7%, IQR 12.7-27.5%) with a low frequency of CD4^+^ (2.7% IQR 0.9-3.4%) MAIT cells identified. This disagrees with the earliest report of MAIT cells in cattle suggesting that cattle MAIT cells were not CD8 positive (20). Interestingly, some of the tetramer positive cells were TCRδ^+^ (mean 8.33% IQR 2.3-13.3% of total MR1-5-OP-RU tetramer^+^ population in PBMC) (**Fig. 2A**), equating to a mean frequency of 0.08% of total γδ^+^ T cells in cattle. Cattle are a γδ T cell high species and in adult cattle ∼10-20% of circulating lymphocytes are TCRγδ positive (50). This observation mirrors a recent report of human MR1 reactive γδ T cells (51).

**Figure 2.**
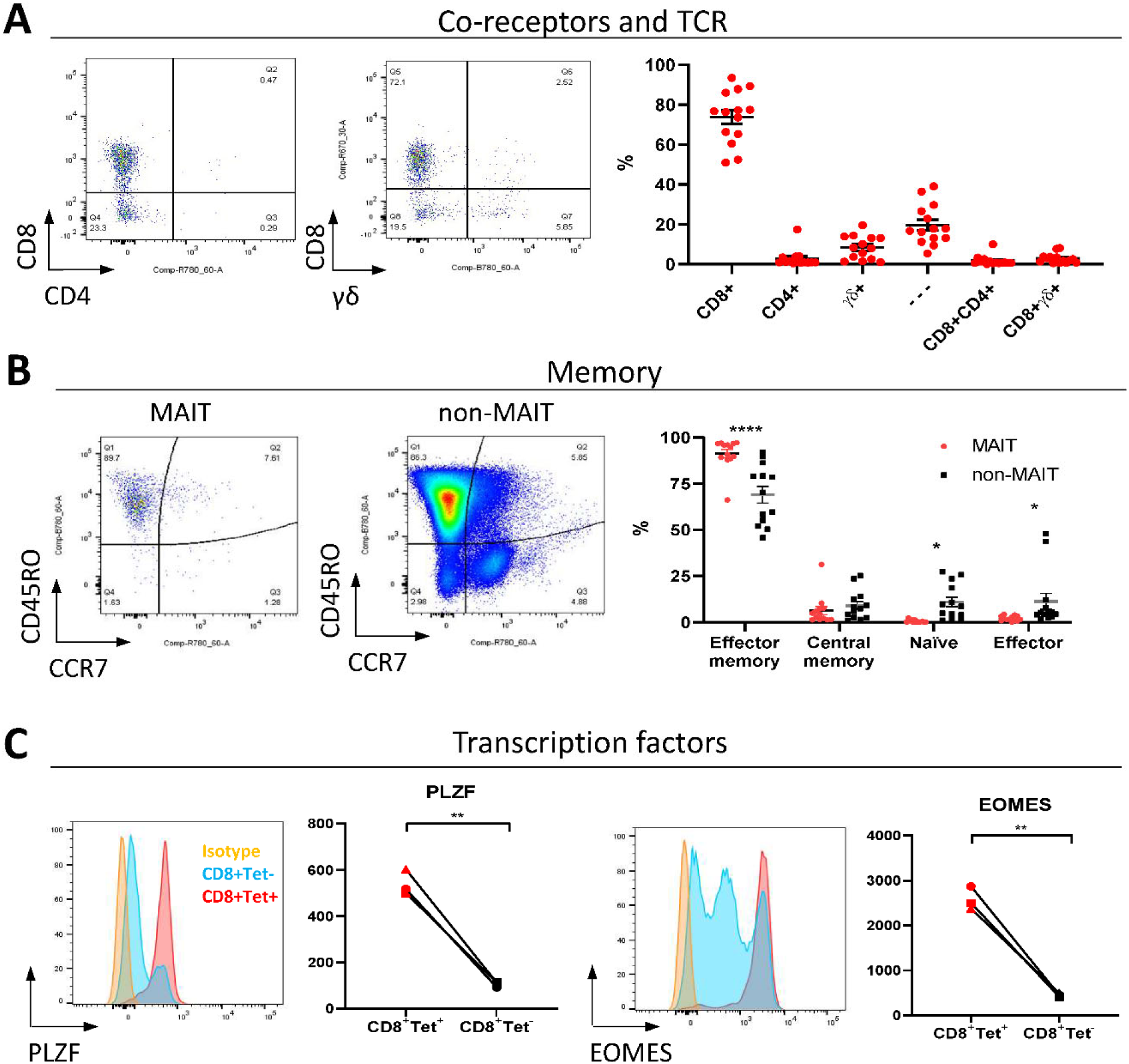
Phenotypic characterisation of bovine MAIT cells in peripheral blood. **(A)** Frequencies of co-receptor and δ-TCR expression of CD3^+^MAIT cells (CD3^+^ MR1-5-OP-RU tetramer^+^) in PBMC. Representative flow cytometry dot plots and summarized data (mean ± SEM, n = 14). CD4^-^CD8^-^δ^-^ = - --. **(B)** Memory phenotyping of peripheral blood MAIT and non-MAIT (CD3^+^ MR1-5-OP-RU tetramer^-^) T cells based on CD45RO and CCR7 expression. Effector memory (CD45RO^+^CCR7^-^), central memory (CD45RO^+^ CCR7^+^), naïve (CD45RO^-^ CCR7^+^), effector (CD45RO^-^ CCR7^-^) phenotype. Data was analyzed using two-way ANOVA with T cell population as repeated measures, followed by Sidak’s multiple comparisons post-hoc test comparing memory phenotype between populations. Representative flow cytometry plots and summarized data (mean ± SEM, n = 13) are shown. **(C)** Expression of the transcription factors PLZF and EOMES in CD8^+^ MR1-5-OP-RU tetramer^+^ MAIT cells (CD8^+^Tet^+^) and non-MAIT T cells (CD8^+^Tet^-^) from PBMC; representative flow cytometry histograms and summarized data of mean fluorescence intensities (n = 3) are displayed. Paired T test within animal.

In contrast to non-MAIT T cells, peripheral blood MAIT cells featured almost exclusively an effector memory phenotype (CD45RO^+^ CCR7^-^) (**Fig. 2B**), as in humans (48). We also compared transcription factor expression in CD8^+^ MAIT cells to CD8^+^ non-MAIT cells. Whilst cattle specific antibodies against the transcription factors PLZF, EOMES, RORγT and T-bet are not available, transcription factors are highly conserved between species and monoclonal antibodies have previously been shown to cross react between species. RORγT and T-Bet showed little expression above isotype control likely due to insufficient cross reactivity. However, PLZF and EOMES-expression was significantly higher in bovine CD8^+^ MR1-5-OP-RU tetramer^+^ MAIT cells compared to CD8^+^ non-MAIT T cells (**Fig. 2C**). Similarly, antibodies specific for human TRAV1-2 (clone 3C10) and CD161 (clone 191B8) did not appear to cross react (data not shown). Together these data showed that MAIT cells in cattle almost exclusively had an effector memory phenotype and were predominantly CD8, PLZF and EOMES positive.

### 3.3 MAIT cells in cattle can be activated by 5-OP-RU and by cytokines

As the majority of MAIT cells were CD8^+^ (**Fig. 2A**), activation experiments focussed on comparing human MR1-5-OP-RU tetramer positive and negative CD8^+^ populations (MAIT and non-MAIT CD8 T cells, respectively). CD8^+^ MAIT cells were specifically stimulated to express IFN-γ (mean 32% IFN-γ^+^) and TNF (mean 29% TNF^+^) by the canonical MAIT cell ligand 5-OP-RU (**Fig. 3A**) at concentrations as low as 50 pM (**Fig. S2A-B)**. Increased concentrations of 5-OP-RU (**Fig. S2C**) and prolonged incubation time with 5-OP-RU (**Fig S2D**) correlated with declining fractions of MR1-5-OP-RU tetramer^+^ cells (**Fig. S2C**), suggesting that the TCRs of bovine MAIT cells are downregulated upon binding to cognate ligand, as described previously (52). The residual fraction of cytokine positive CD8^+^ tetramer negative cells following 5-OP-RU stimulation (**Fig 3A, S2**) are therefore most likely activated MAIT cells with downregulated TCRs. The 5-OP-RU-mediated activation of cattle MAIT cells was competitively inhibited by the inhibitory MR1 ligand Ac-6-FP (**Fig. 3B**), as is the case with human (41) and mouse MAIT cells (53), and strongly suggests that activation is mediated through MR1-TCR interactions.

**Figure 3.**
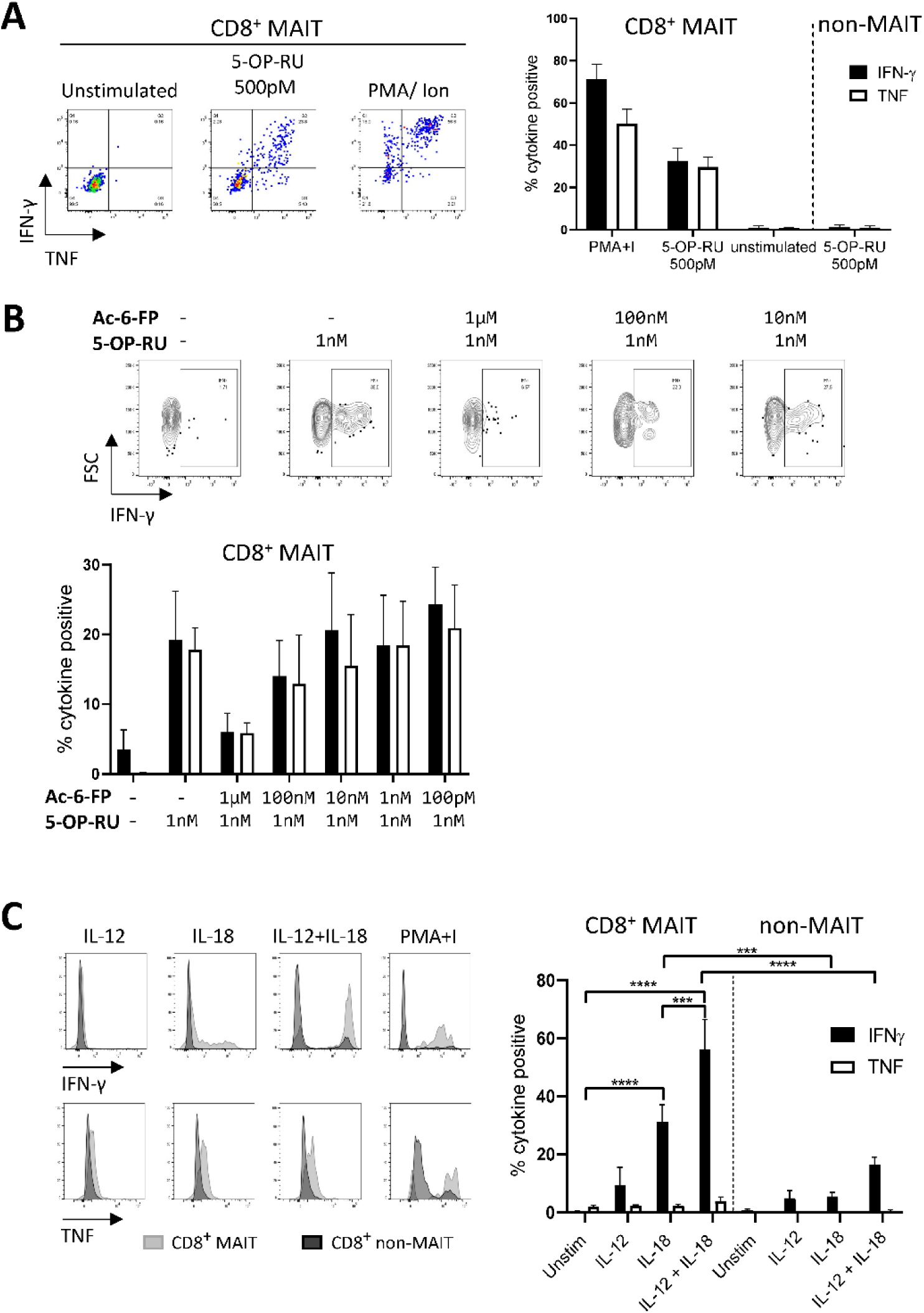
Antigen and cytokine stimulation of cattle MAIT cells. **(A)** Representative flow cytometry plots and summarized data of IFN-γ and TNF expression (mean ± SEM, n = 3) of live CD8^+^ MR1-5-OP-RU tetramer^+^ (MAIT) and CD8^+^ tetramer^-^ (non-MAIT) cells incubated with 5-OP-RU for 7 h at a concentration of 500 pM. Unstimulated cells and phorbol myristate acetate & ionomycin (PMA+I) stimulated controls are also shown. **(B)** Representative flow cytometry plots and summarized data of cattle PBMC incubated with Ac-6-FP prior to the addition of 1 nM 5-OP-RU (mean ± SEM, n = 3). Lymphocytes were gated for FSC&SSC, singlets, live and CD8 expression followed by tetramer^+^ / IFN-γ^+^ gating. The fraction of IFN-γ and TNF positive cells within MAIT cells (CD8^+^ tetramer^+^ or IFN-γ^+^) was determined. **(C)** PBMC were stimulated for 18 hours with IL-12, IL-18 or a combination of both (all at 50 ng/ml). Representative flow cytometry histogram plots show IFN-γ and TNF expression within CD8^+^ MAIT cells overlaid with CD8^+^ non-MAIT cells. Summarized data of the fraction of cytokine positive CD8^+^ MAIT and non-MAIT cells are shown (mean ± SEM, n = 4). Data were analyzed using two-way ANOVA with repeated measures within animal, followed by Sidak’s multiple comparisons post-hoc test comparing the mean of each cytokine stimulation within and between CD8^+^ MAIT cells and CD8^+^ non-MAIT cells.

Viruses can stimulate MAIT cells in a TCR independent manner via cytokine stimulation (54). Following stimulation with IL-18, bovine MAIT cells produced IFN-γ (mean 31 % IFN-γ^+^) and low frequencies of TNF producing cells (mean 2 % TNF^+^) were observed (**Fig. 3C**). There was no significant response to stimulation with IL-12 only by MAIT or non-MAIT T cells, but in conjunction with IL-18, IL-12 did significantly increase the frequency of MAIT cells producing IFN-γ (mean 56% IFN-γ^+^) compared to IL-18 stimulation alone. Unlike 5-OP-RU stimulation, cytokine stimulation did not appear to affect MR1-5-OP-RU tetramer binding, suggesting that it did not induce TCR downregulation (data not shown).

### 3.4 Cattle MAIT cells express a conserved T cell receptor alpha chain and show low beta chain diversity

The canonical MAIT TCRα chain has previously been identified in cattle by sequencing of a limited number of unsorted T cells with no pre-identification of MAIT cells (20, 23). Here we performed deep TCR profiling of sorted bovine CD8^+^ MR1-5-OP-RU tetramer^+^ MAIT cells in comparison to MR1-5-OP-RU tetramer^-^ CD8^+^ T cells. The sorted MAIT cell populations were highly enriched for the canonical *TRAV1* (73.3%) and *TRAJ33* (72%) gene segments which were not enriched in the CD8^+^ non-MAIT cell population (**Figs. 4A-B, S3 and S4**). The TCRβ chain usage of bovine MAIT cells was more diverse than the TCRα chain, though an enrichment of *TRBV4, TRBV7 and TRBV20* was seen in the MAIT cell population compared to the non-MAIT CD8^+^ population (**Figs. 4A-B, S3 and S4**), accounting for 35 %, 13 % and 23 % of total TRBV sequences in MAIT cells respectively. This mirrors findings in humans where the critical residues for MR1 recognition are found in the TCRα chain (5, 55) and the TCRβ chains are more variable, though specific TRBV, particularly *TRBV6 and TRBV20*, dominate (19, 21, 38, 41, 56, 57). The nomenclature of cattle TCR gene segments is based on human orthologues and *TRBV20* is, therefore, enriched in both human and cattle MAIT cells. The CDR3α loops of CD8^+^ MAIT TCRs were primarily 12aa long, similar to the canonical human MAIT cell TCR (**Fig. 4C**), accounting for 76 % of all MAIT cells, whereas CDR3α loops of non-MAIT TCRs varied much more in length and most (73 %) were longer. The CDR3β loops of CD8^+^ MAIT TCRs were more varied in length than the CDR3α loops and displayed a similar length distribution as compared to those of non-MAIT CD8^+^ TCRs. In humans the MAIT TCRα rearrangements *TRAV1-2-TRAJ33/20/12* account for the majority (∼95%) of MAIT TCR clonotypes in blood (19, 21, 38, 56, 58).

**Figure 4.**
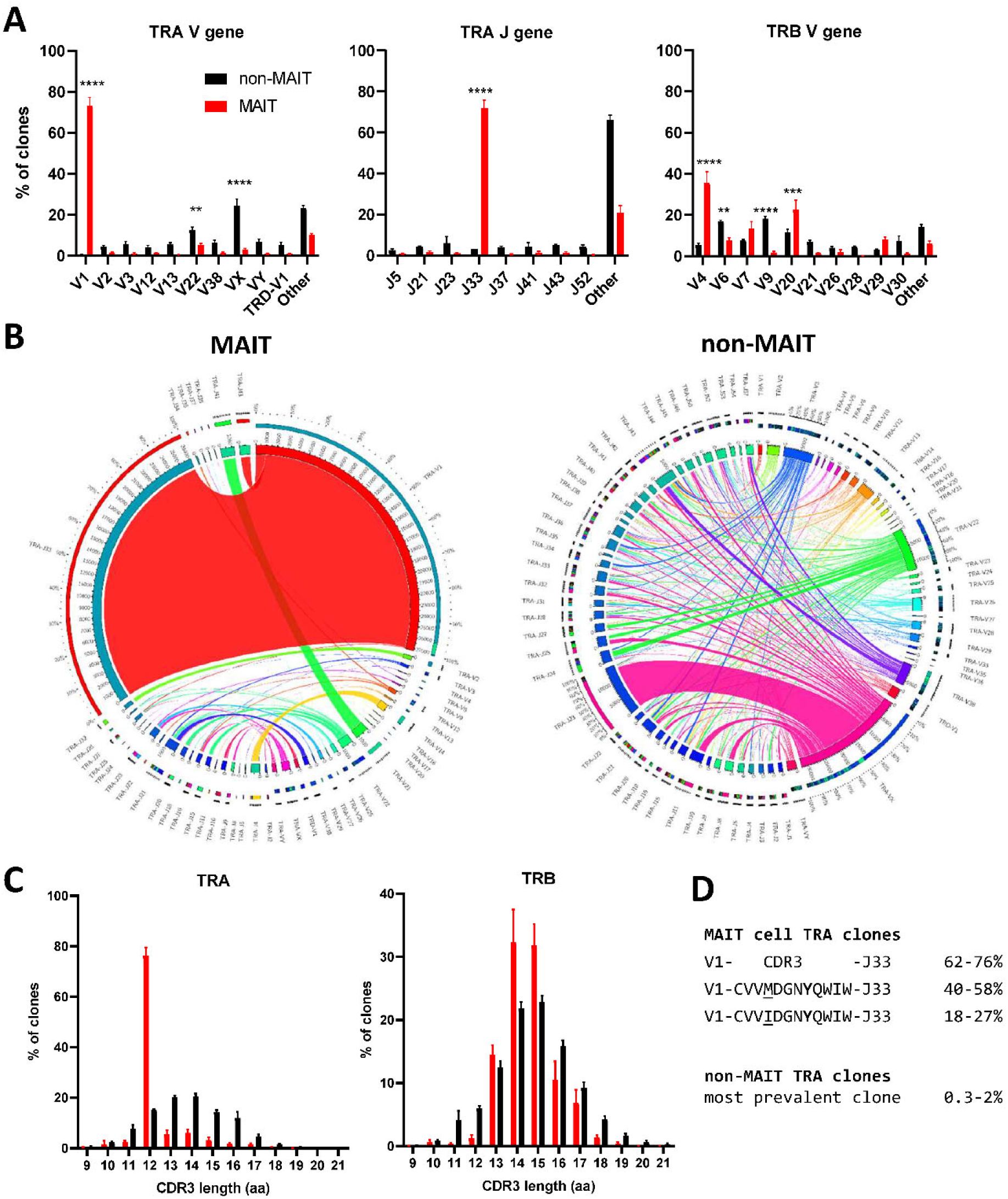
Conserved TCR alpha chain and limited beta chain diversity in bovine MAIT cells. **(A)** Bar graphs showing TCR alpha (TRA) V and J and TCR beta (TRB) V gene usage in MAIT (CD8^+^ MR1-5-OP-RU tetramer^+^) and non-MAIT (CD8^+^ MR1-5-OP-RU tetramer^-^) T cells. Bars represent the percentage of total TRA or TRB sequences obtained from high throughput TCR sequencing of populations sorted from PBMC (mean ± SEM, n = 4, number of sequences in Supplementary table 1). Data were analyzed using two-way ANOVA with repeated measures within animal, followed by Sidak’s multiple comparisons post-hoc test comparing gene usage between MAIT and non-MAIT T cells. **(B)** Circos plots showing TRAV-J combinatorial diversity within MAIT and non-MAIT T cells. The inner circle shows the number of sequence reads per V and J gene and TRAV/J combinations are indicated by proportional bands linking the genes using the colour of the V gene. The outer ring shows the frequency of pairing for each TCR gene segment to the reciprocal gene segment coloured according to the paired gene. Representative plots from a single individual. **(C)** Distribution of TRA and TRB CDR3 amino acid (aa) length. **(D)** Alignment of the TCR alpha CDR3 region of the dominant MAIT cell clones.

When analyzing the TCR sequences at the clonal level, 62-76% of CD8^+^ MAIT cells expressed the canonical *TRAV1*-*TRAJ33* TCR rearrangement (**Fig. 3B, S3 and S4**), similar to what Greene et al. (25) previously described in macaques where 70% of TCRs of MR1 tetramer positive peripheral blood T cells were *TRAV1-2*^+^. The non-canonical TCR rearrangements were highly diverse, with no CDR3 sequences shared between all four donors (**Supplementary Data 1**). The non-canonical TCR may be the result of nonspecific binding of tetramer to non-MAIT cells, sorting impurities or non-MAIT cell MR1 reactive T cells, which are rare populations identified in mice and humans (59). *TRAV1*-*TRAJ33*^+^ cattle MAIT TCRs featured two similar CDR3 sequences, the predominant sequence of which was CVVMDGNYQWIW with a secondary sequence observed in all animals with a single aa substitution of methionine to isoleucine at position 91 (**Fig. 3D, Supplementary Data 1**). The same position in addition to the neighbouring residue also varies in human *TRAV1-2*-*TRAJ33*^+^ MAIT cell TCRs (CAXXDSNYQLIW)(20, 38, 57, 58). Both of the cattle CDR3α sequences are conserved in Tyr95, which is critical for MR1 and antigen binding in humans (55). While cattle have orthologues for the human *TRAJ12* and *TRAJ20* gene segments, including conservation of a tyrosine at the same position, these *TRAJ* segments were not enriched in cattle MAIT cells. Altogether, the identified TCR CDR3α sequences are in agreement with previously predicted putative cattle MAIT TCR CDR3α sequences (20, 23).The deep TCR profiling, together with the functional and phenotypical analyses, confirmed that the MR1-5-OP-RU tetramer positive cells in cattle are bona fide MAIT cells.

### 3.5 Distribution and phenotypic comparison of MAIT cells in tissues

In cattle PBMC, approximately 0.6% of CD3^+^ lymphocytes (**Fig. 1B**) or 4% of CD8^+^ lymphocytes were MAIT cells (**Fig. 5A-B**). MAIT cells in humans are highly enriched in mucosal tissues and liver (2). This was also true in cattle (**Fig. 5A-B**), with greater frequencies of MR1-5-OP-RU tetramer positive cells detected in lung, spleen, liver and BAL when compared to PBMC (mean 1.74%, 1.02%, 2.71% and 1.84% of CD3^+^ lymphocytes respectively). MAIT cell frequencies in the ileum were comparable to those in PBMC (mean 0.6%). Of note, an enrichment of MAIT cells was observed in the mesenteric Ln (mean 3.5%), but not in the pre-scapular Ln (mean 0.6%) (**Fig. 5A-B**). The high frequency of MAIT cells in the mesenteric Ln was not seen in a recent study of the pigtail macaque (39) and may be specific to cattle. One could speculate that this difference is due to the large microbial populations in the rumen and large intestines of cattle, which are drained by the mesenteric lymph node. Although in all tissues a low fraction of T cells bound MR1-6-FP tetramer compared to MR1-5-OP-RU tetramer, relatively more MR1-6-FP tetramer positive T cells were identified in spleen and liver (**Fig. 5A-B**). The percentage of effector memory non-MAIT T cells varied greatly across tissues, whereas MAIT cells predominantly had an effector memory phenotype irrespective of origin (**Fig. 5C**). Non-MAIT T cells in lymph nodes were predominantly CCR7 high. In contrast, MAIT cells in prescapular and mesenteric lymph nodes had an effector memory phenotype with low expression of CCR7, as was reported for human MAIT cells in thoracic duct lymph (60). Migration of γδ T cells from tissue to lymph nodes was shown to be CCR7 independent in cattle (61) and it has been hypothesized that CCR7 low MAIT cells enter the lymphatics from tissues in a CCR7 independent manner (60).

**Figure 5.**
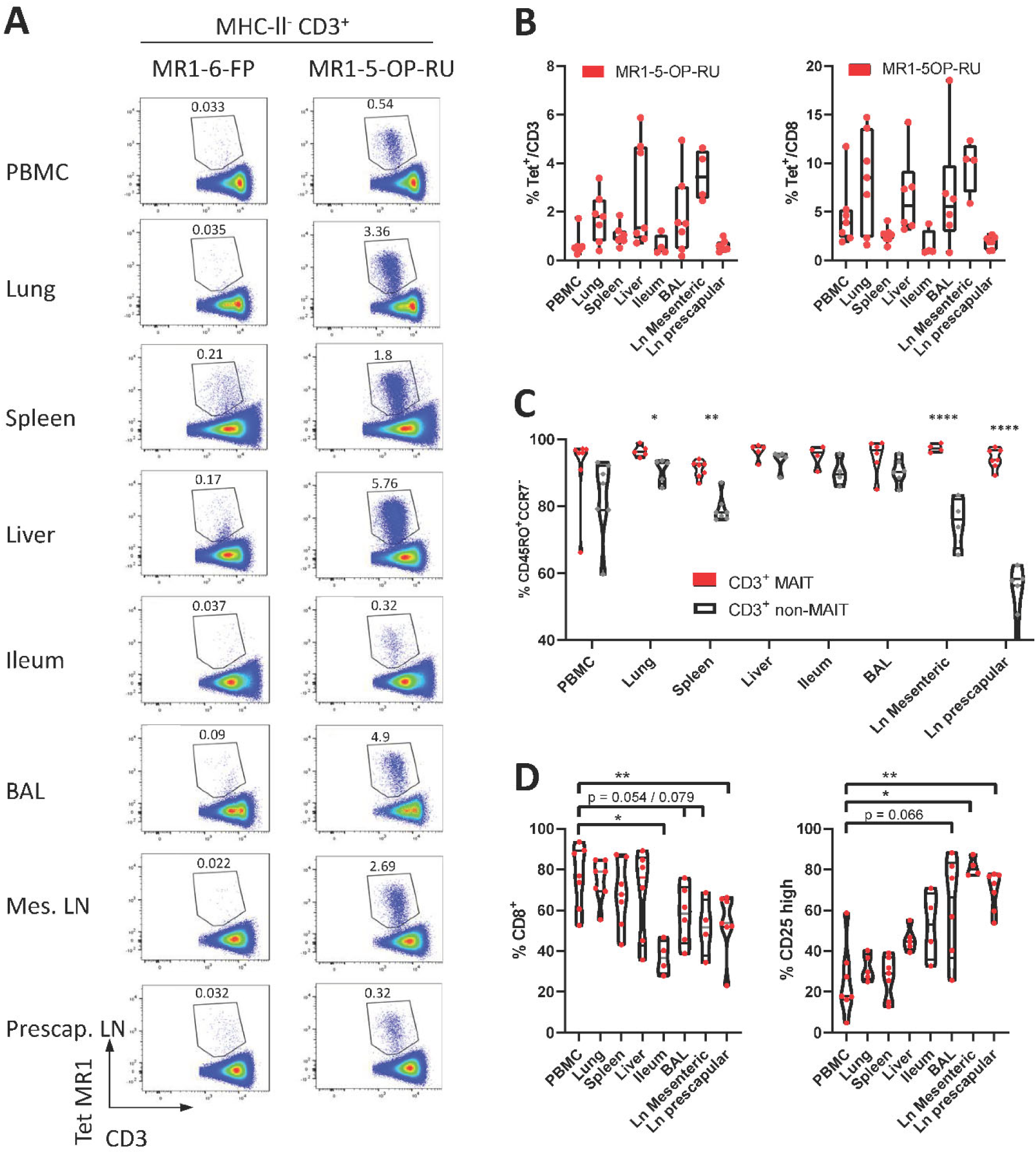
Comparisons of the frequencies and phenotype of cattle MAIT cells in different tissues. Representative flow cytometry plots of MR1-5-OP-RU tetramer gating within MHC-II^-^ CD3^+^ cells in different tissues. **(B)** Frequencies of CD3^+^ MR1-5-OP-RU tetramer^+^ MAIT cells and CD8^+^ MR1-5-OP-RU tetramer^+^ MAIT cells in different tissues from cattle. Number of animals per tissue are given in supplementary table 2. **(C)** Frequency of effector memory (CD45RO^+^CCR7^-^) MAIT (CD3^+^ tetramer^+^) and non-MAIT T cells (CD3^+^ tetramer^-^). Mixed effects model with matching across tissue and cell population, followed by Sidak’s multiple comparisons post-hoc test comparing differences between MAIT and non-MAIT cells within tissues. **(D)** Frequency of CD8^+^ or CD25 high MAIT cells (CD3^+^ MR1-5-OP-RU^+^). Mixed effects model with matching across tissue, followed by Dunnet’s multiple comparisons post-hoc test comparing differences between PBMC and other tissues.

Differences in MAIT cell co-receptor usage between blood and tissues were observed (**Fig. 5D**). MAIT cells from peripheral blood had the largest CD8^+^ population (mean 76% CD8^+^) with no significant difference in the frequency of MAIT cells expressing CD8 in lung, spleen and liver (mean 74%, 68% and 68% CD8^+^ respectively). Significantly lower fractions of CD8^+^ MAIT cells were identified in the pre-scapular Ln. (mean 54% CD8^+^) and in ileum, which showed the lowest frequency of CD8 expression (mean 37% CD8^+^). There was a trend for a lower fraction of CD8^+^ MAIT cells in mesenteric Ln. (mean 52% CD8^+^, p = 0.079) and in BAL (mean 58%, p = 0.054). There was a strong negative correlation between the fraction of CD8^+^ and CD8^-^CD4^-^TCRγ^-^ MAIT cells (R^2^ = 0.90) and the fraction of these triple negative MAIT cells was proportionally increased in tissues with low CD8 expression. Differences in IL-2Rα chain (CD25) expression were also noted (**Fig. 5D**), with a trend for a greater frequency of CD25 high MAIT cells in BAL (mean 61%, p = 0.066), compared to MAIT cells in peripheral blood (mean 25%). A significantly higher proportion of CD25 high MAIT cells was seen in prescapular (mean 69%) and mesenteric (mean 81.2%) lymph nodes, which is more comparable to other tissues than to blood, potentially due to MAIT cell recirculation between tissues and lymph nodes (60, 61).

### 3.6 Cattle MAIT cells respond to bacterial infections *in vivo* and bacterial stimulation *in vitro*

Next, we sought to characterise MAIT cells in cattle directly *ex vivo* during infection as well as in an immunisation-challenge model. Mastitis is an inflammation of the mammary gland and is most often due to bacterial infections by riboflavin biosynthesis proficient bacteria, such as *Escherichia coli* (*E. coli*) and *Staphylococcus aureus* (*S. aureus*). It is the most frequent disease in dairy cattle, presents a major impact on animal welfare, and is associated with economic losses (62). Milk contains many different blood derived immune cells (63, 64), and MR1-tetramer staining of cells in milk from healthy cows revealed a distinct MAIT cell population consistently present in bovine milk (mean 0.8% of CD3^+^, IQR 0.4-1.4%, n = 6, **Fig 6A, S5A**). The number of cells in milk, also referred to as somatic cell count (SCC) is used as a biomarker for mastitis, where animals with an elevated SCC (>200,000 cells/ml) are considered to have mastitis (65). Cattle with an elevated SCC had on average a greater than 5-fold increase in MAIT cells as a percentage of CD3^+^ T cells, indicating increased migration of MAIT cells relative to other T cells from blood to milk during mastitis (**Fig. 6A**) and suggesting a possible direct or bystander role of MAIT cells in this inflammatory condition.

**Figure 6.**
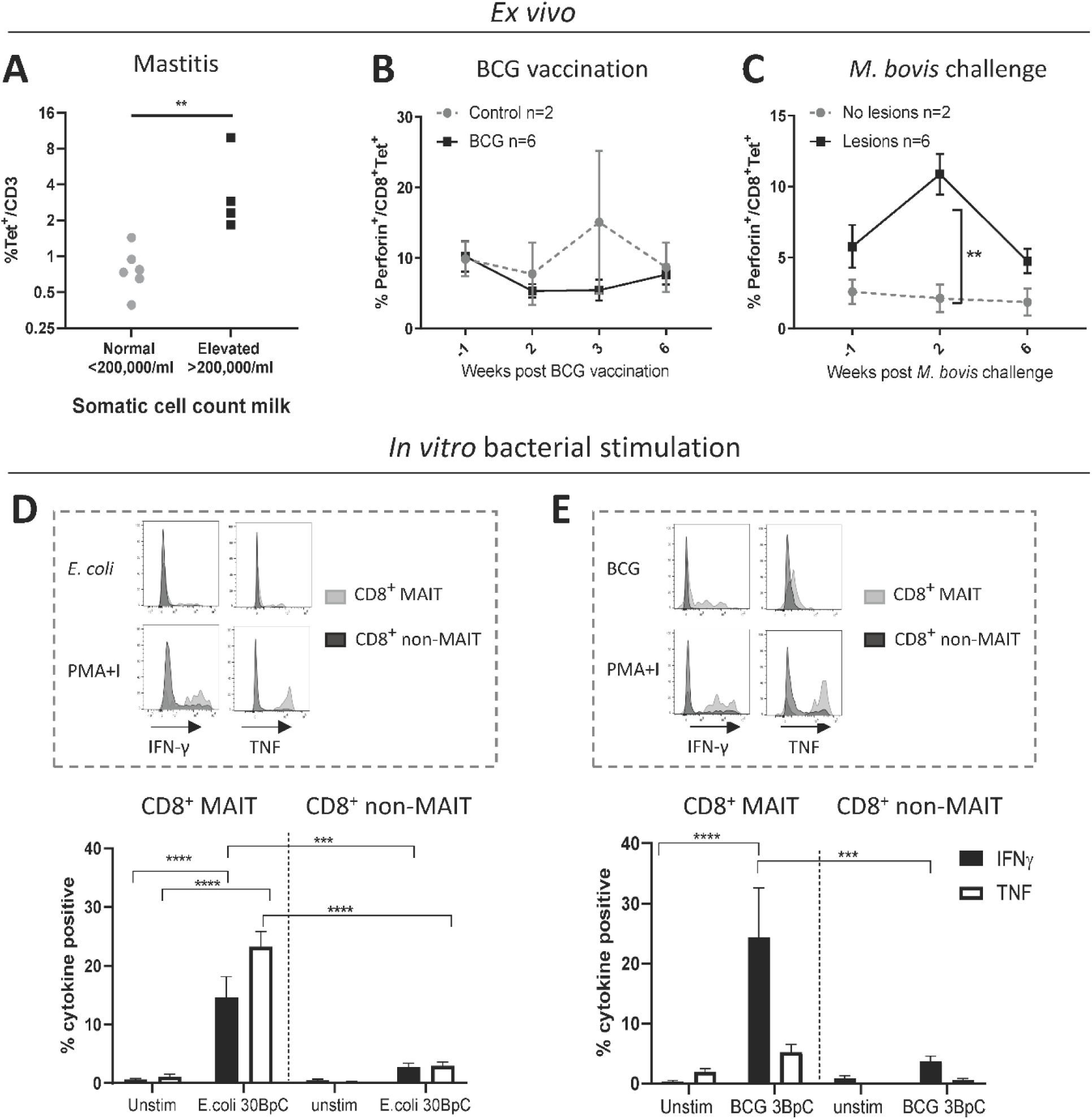
Cattle MAIT cells respond to bacterial infections *in vivo* and *in vitro*. **(A)** The percentage of MAIT cells (MR1-5-OP-RU tetramer^+^) within the total T cell population (CD3^+^) in milk, grouped according to normal (<200.000 cells/ml) versus elevated milk somatic cell count (SCC) (>200.000 cells/ml, indicating a mastitis). Groups were compared using a Mann–Whitney U test. **(B)** Six calves were vaccinated with 4.6 × 10^6^ CFU BCG Danish SSI 1331 subcutaneously and activation of MAIT cells was measured longitudinally as the percentage of perforin^+^ CD8^+^ MR1-5-OP-RU tetramer^+^ MAIT cells in PBMC. **(C)** MAIT cell activation was measured longitudinally, as in B, following *Mycobacterium bovis* challenge. Calves (n = 8) were challenged endobronchially with 10^4^ CFU *M. bovis* AF2122/97. Animals were grouped based on the presence of tuberculosis associated lesions in lungs and lymph nodes at post-mortem examination 11 weeks post-challenge. **(B**,**C)** Data were analyzed using two-way ANOVA with time as repeated measures, followed by Sidak’s multiple comparisons post-hoc test comparing differences between groups at each time point. Mean ± SEM is indicated. **(D and E)** Cattle PBMC were stimulated for 7 hours with 30 *E. coli* bacteria per cell (BpC) (**D**) or for 18 hours with 3 *M. bovis* BCG BpC (**E**). Representative flow cytometry histograms and summarized data of IFN-γ and TNF expression in live CD8^+^ MR1-5-OP-RU tetramer^+^ MAIT and CD8^+^ MR1-5-OP-RU tetramer^-^ non-MAIT cells are shown (mean ± SEM, n = 4). Data were analysed using two-way ANOVA with repeated measures within animal, followed by Sidak’s multiple comparisons post-hoc test comparing IFNγ and TNF expression of stimulated CD8^+^ MAIT cells to unstimulated MAIT cells and to non-MAIT cells.

We next assessed CD8^+^ MAIT cell responses to *Mycobacterium bovis* (*M. bovis*) infection longitudinally in cattle vaccinated with the attenuated *M. bovis* strain Bacillus Calmette–Guérin (BCG) and following endobronchial challenge with the virulent *M. bovis* strain AF2122/97. Perforin and granzyme production can be used as activation markers of MAIT cells (2, 66). In cattle, no changes in the frequency of perforin expressing MAIT cells were found *ex vivo* in PBMC following BCG vaccination (**Fig. 6B**). However, two weeks following endobronchial challenge with *M. bovis* the fraction of perforin expressing CD8^+^ MAIT cells was significantly higher in animals that showed tuberculosis associated lesions in the lungs and lymph nodes compared to animals without lesions (**Fig. 6C**). While perforin expression did not change in tetramer negative (non-MAIT) CD8^+^ T cells, there was a significant, transient, increase in perforin expression amongst CD8^+^ MAIT cells in animals with lesions (**Fig. S5B**). In macaques, activation of MAIT cells was much more pronounced locally at the site of BCG vaccination (25). Vaccination with the attenuated *M. bovis* BCG strain causes a local infection and it is therefore not surprising that MAIT cell activation was not detectable in peripheral blood. We hypothesize that severe infection with virulent *M. bovis* resulting in lesions in multiple organs leads to more widespread MAIT cell activation that can be detected in peripheral blood. No changes in CD69 expression, or in CD8^+^ MAIT cell frequencies were detected in peripheral blood of BCG vaccinated or *M. bovis* challenged animals (**Fig. S5C-F**), which is in agreement with findings after BCG vaccination in humans (67) and *M. tuberculosis* challenge in macaques (25). Overall, these data demonstrate that *M. bovis* infection in cattle can lead to activation of MAIT cells *in vivo*.

Having established that MAIT cells may respond to bacterial infections in cattle *in vivo*, we went on to validate whether cattle MAIT cells were activated by riboflavin biosynthesis competent bacteria. PBMC were stimulated with *E. coli*, and the attenuated *M. bovis* strain BCG (**Figs. 6D-E**). Stimulation with *E. coli* for 7 hours led to robust IFN-γ and TNF upregulation by MAIT cells, while tetramer negative CD8^+^ T cells showed limited activation **(Fig. 6D)**, which may include activated MAIT cells that have downregulated their TCRs (**Fig. S2**). When stimulated overnight with BCG, MAIT cells displayed robust IFN-γ production whilst TNF expression was limited (**Fig. 6E**), comparable with the cytokine profile observed upon IL-12/IL-18-stimulation (**Fig. 3C**). This is in agreement with human MAIT cell responses to BCG stimulation, which were reported to be mediated primarily by IL-12/IL-18 rather than TCR-antigen-MR1 stimulation and yielded INF-γ, but not TNF production (67). *S. aureus* also stimulated IFN-γ and TNF production in bovine MAIT cells (data not shown). Together these data illustrate that cattle MAIT cells respond to bacterial infections *in vivo* and are activated by bacteria *in vitro*.

## 4 Discussion

The canonical MAIT cell TCR α chain was first identified in cattle alongside humans and mice over 20 years ago (20), but phenotypic and functional MAIT cells have not been described in any livestock species. The use of human MR1 tetramers that cross react with cattle have allowed us to identify MAIT cells in cattle and characterise their phenotype and function *in vitro* and directly *ex vivo*. While these data were generated using a xeno-MR1 reagent, the further phenotypic and functional analysis of cattle MAIT cells was in great agreement with that of other species and thus gives confidence that the human MR1-5-OP-RU tetramer identifies a MAIT cell population in cattle. Our data demonstrate that cattle MAIT cells are phenotypically and functionally similar to their human counterparts, including expression of an orthologous conserved TRAV1-TRAJ33 T cell receptor α chain by the majority of MR1 tetramer^+^ cells, an effector memory phenotype, expression of transcription factors associated with innate immunity, enrichment in mucosal tissues and activation by synthetic 5-OP-RU, the cytokines IL-12 and IL18, and riboflavin biosynthesis competent bacteria.

Cattle produce around 20 to 60 litres of milk per day and are milked at least twice a day. The large volumes and continuous production of milk means there is a huge migration of immune cells from blood to milk, even in a healthy non-infected non-inflamed mammary gland (63, 64). The increased MAIT cell frequency in milk in cows with mastitis implies increased trafficking of MAIT cells relative to other T cells to the mammary gland during infection. Mastitis in cattle is predominantly bacterial in origin and is characterised by a massive migration of neutrophils to the mammary gland (68). The major mastitis pathogens *E. coli* and *S. aureus* (69) stimulated bovine MAIT cells *in vitro*. MAIT cells are a key source of pro-inflammatory cytokines (30, 37, 70) and bacteria induced cytokine responses by MAIT cells in the context of mastitis could be a driving force in the neutrophil influx and inflammation associated with intramammary bacterial infections. Further studies tracking MAIT cells longitudinally in milk and tissues during intramammary infections, will shed light on the role of MAIT cells in mastitis, including the temporal relation to neutrophil influx. Maternal immune cells in milk play a role in the development of the neonatal immune system (71, 72) and milk derived CD8^+^ T cells preferentially home to the payers patches of the small intestine (73). MAIT cells have also been identified in human breast milk (74). Given the monomorphic nature of their restriction element MR1, MAIT cells are donor-unrestricted and can be activated by MR1 expressing cells from any individual (75). MAIT cells present in milk and possibly also in colostrum may be transferred to the neonate where they could play a role in passive immunity in the upper and lower intestinal tract (72).

MAIT cells have been shown to be activated by *Mycobacterium tuberculosis* in humans (13, 67), non-human primates (40) and mice (76). Furthermore, MAIT cells are the predominant IFN-γ producing T cell population in TB exposed individuals upon restimulation with BCG (67). The increased proportion of perforin^+^ MAIT cells in cows with TB lesions combined with the activation of MAIT cells by BCG *in vitro* indicates that MAIT cells may play a role in bovine TB. Intravenous, but not intradermal, administration of BCG was shown to transiently (up to 8 weeks) increase MAIT cell frequencies in the BAL of non-human primates, while there was no effect on MAIT cells in peripheral blood (77). The same study also showed limited responses by MAIT cells in the periphery to subcutaneous BCG vaccination and demonstrates that route of vaccine administration and tissue localisation are important factors to consider when studying MAIT cell responses to vaccination. Recently, 5-OP-RU vaccination of mice was not shown to be protective of TB infection and contributed to a delayed CD4 response to the infection. However, treatment with 5-OP-RU during chronic TB infection led to an increase in MAIT cell frequencies and a lowering of bacterial burden, which was dependent on IL-17 expression (78). The emerging picture suggests that MAIT cells are involved in immunity against TB infection, although whether their role is protective may well depend on a range of factors, including stage of infection. As a natural host of TB and with the possibility for repeated (tissue) sampling and cannulation of lymph nodes, cattle are an appropriate large animal model with unique potential to study MAIT cells longitudinally in tissues *in vivo*.

In summary, we have identified a MAIT cell population in cattle with phenotypic and functional characteristics closely resembling MAIT cells in mice and humans. We have demonstrated that cattle MAIT cells respond to bacterial infections of economic and zoonotic importance and the data and tools presented here will facilitate the use of cattle as a relevant large animal model to study MAIT cell biology during immunisation and infection.

## 5 Acknowledgements

We are grateful to the staff of the Dryden farm of the Roslin Institute and the animal staff at the APHA for their invaluable help with the animal experiments. We also thank the Pirbright flow cytometry facility (National capability science services) for their support with flow cytometry and the Roslin Institute Veterinary Immunological Toolbox facility for support with monoclonal antibody production.

## 7 Author contribution statement

PK, ET, LB, SE, TKC conceived, designed and coordinated the study. ME, LB, TKC, CV, SJ, MV, EM, TC designed and performed experiments, processed samples and analyzed the data. JM, LL, DF generated 5-OP-RU. LB, ME, SE, prepared the manuscript and figures. All authors reviewed the manuscript and approved the submitted version.

## Supplementary Material

### 12 Supplementary tables

**Supplementary table 1.** Number of paired sequence reads after quality control and TCR database alignment for TCR alpha (TRA) and beta chain (TRB) sequencing of MAIT (CD8^+^ MR1-5-OP-RU tetramer^+^) and non-MAIT (CD8^+^ MR1-5-OP-RU tetramer^-^) T cells sorted from PBMC (n = 4).

|  | TRA |  | TRB |  |
| --- | --- | --- | --- | --- |
|  | non-MAIT | MAIT | non-MAIT | MAIT |
| **Cow 1** | 97999 | 68947 | 80861 | 165486 |
| **Cow 2** | 95250 | 34336 | 33350 | 43914 |
| **Cow 3** | 46090 | 31828 | 48989 | 126600 |
| **Cow 4** | 68335 | 46797 | 82317 | 89044 |

**Supplementary table 2.** Number of animals per tissue and staining panel for phenotypic characterisation of MAIT cells (CD8^+^ MR1-5-OP-RU tetramer^+^) in tissues. Ln Mes. – Mesenteric lymph node. Ln Prescap. – Prescapular lymph node.

|  | PBMC | Lung | Spleen | Liver | Ileum | BAL | Ln Mes. | Ln Prescap. |
| --- | --- | --- | --- | --- | --- | --- | --- | --- |
| CD3<sup>+</sup>Tet<sup>+</sup> | 7 | 7 | 7 | 7 | 4 | 7 | 4 | 7 |
| CD8<sup>+</sup>Tet<sup>+</sup> | 7 | 7 | 7 | 6 | 4 | 6 | 4 | 7 |
| % CD45RO<sup>+</sup>CCR7<sup>-</sup> | 7 | 5 | 7 | 5 | 4 | 6 | 4 | 7 |
| % CD25 high | 7 | 5 | 7 | 5 | 4 | 6 | 4 | 7 |
| TCR & Co-receptor | 7 | 7 | 7 | 6 | 4 | 6 | 4 | 7 |

**Supplementary figure 1.**
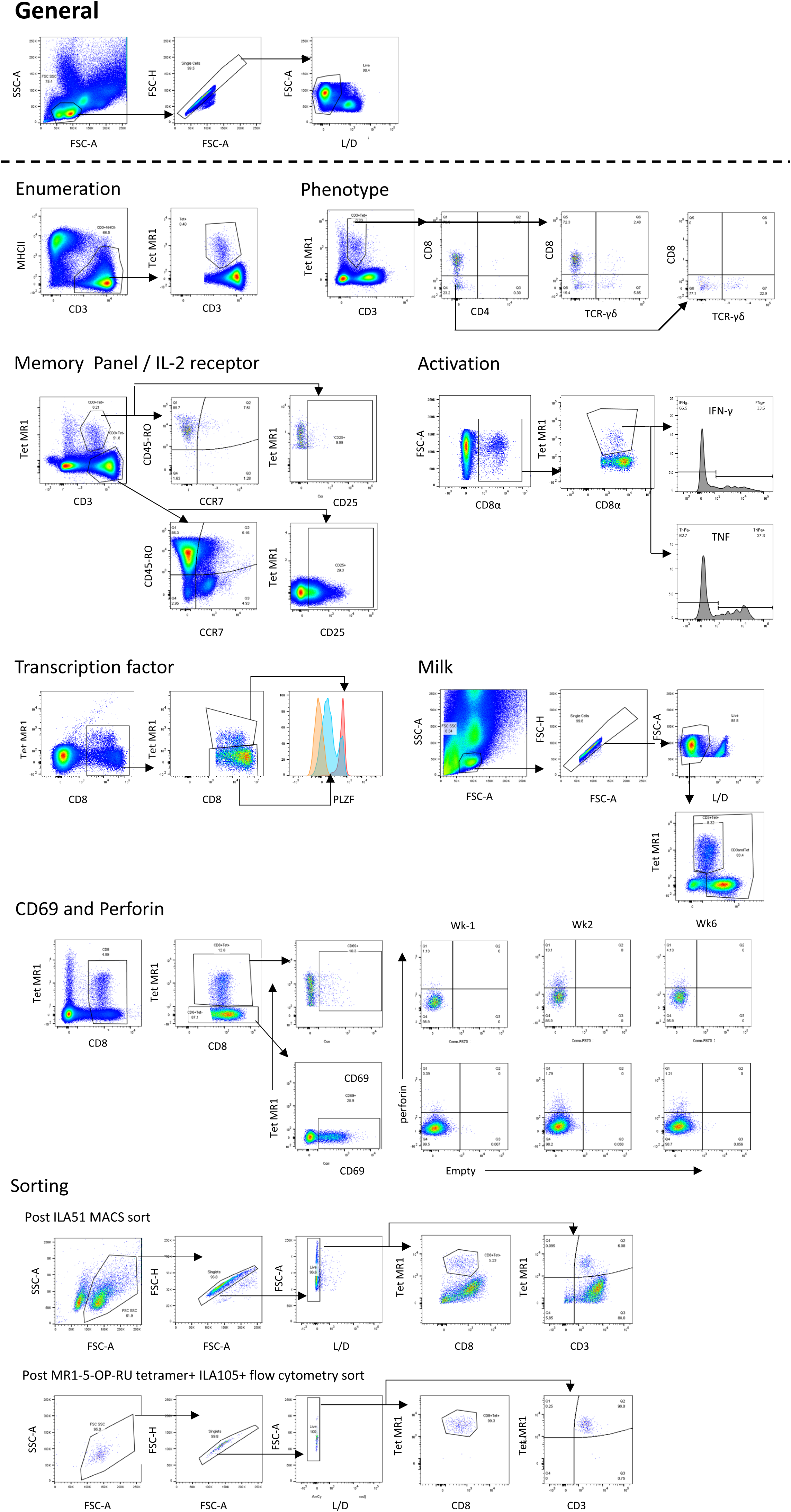
Flow cytometry gating strategy. The general gating strategy was used in all experiments to identify live singlet cells, followed by the specific gating strategies as indicated for each assay / experiment.

**Supplementary figure 2.**
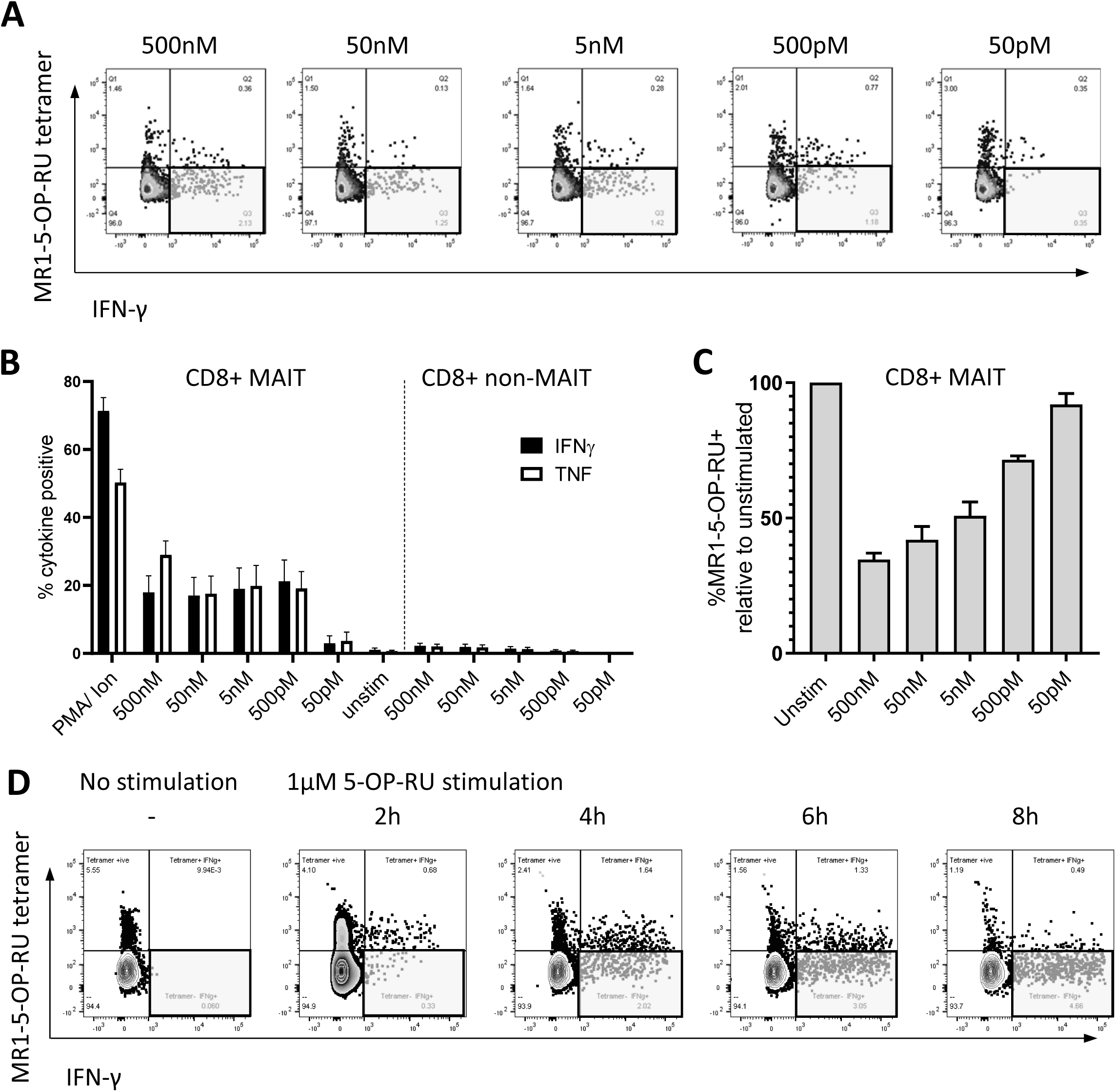
TCR downregulation upon activation of MAIT cells. (**A**) PBMC were stimulated with various concentrations of 5-OP-RU (50 pM – 500 nM) for 7 hours. Lymphocytes were gated on FSC&SSC, singlets, live and CD8 expression. Flow cytometry plots from a representative animal show MR1-5-OP-RU tetramer vs. IFN-γ staining. Grey highlight boxes indicate IFN-γ^+^ T cells which are not detected with MR1-5-OP-RU tetramer. (**B**) Cattle PBMC were stimulated for 7 hours with 5-OP-RU (50 pM – 500 nM), 1μM Ac-6-FP, medium control or PMA/Ionomycin (PMA/Ion). The fractions of IFNγ^+^ or TNF^+^ CD8^+^MR1-5-OP-RU tetramer^+^ MAIT cells and, for comparison, of CD8^+^tetramer^-^ non-MAIT cells are depicted (data combined from two experiments, mean ±SEM, n=6 for each concentration other than 50pM where n = 3). (**C**) The fraction of MR1-5-OP-RU tetramer^+^ MAIT cells following stimulation with 5- OP-RU relative to the fraction of MR1-5-OP-RU tetramer^+^ MAIT cells in unstimulated PBMC from the same animal (mean ± SEM, n=6). (**D**) Bovine PBMC were stimulated with 1 μM 5-OP-RU and 50 ng/ml IL- 12 and IL-18 over a time course of 2 to 8 hours. Flow cytometry gating as in A. Grey highlight boxes indicate IFN-γ^+^ T cells not detected by MR1-5-OP-RU tetramer.

**Supplementary figure 3.**
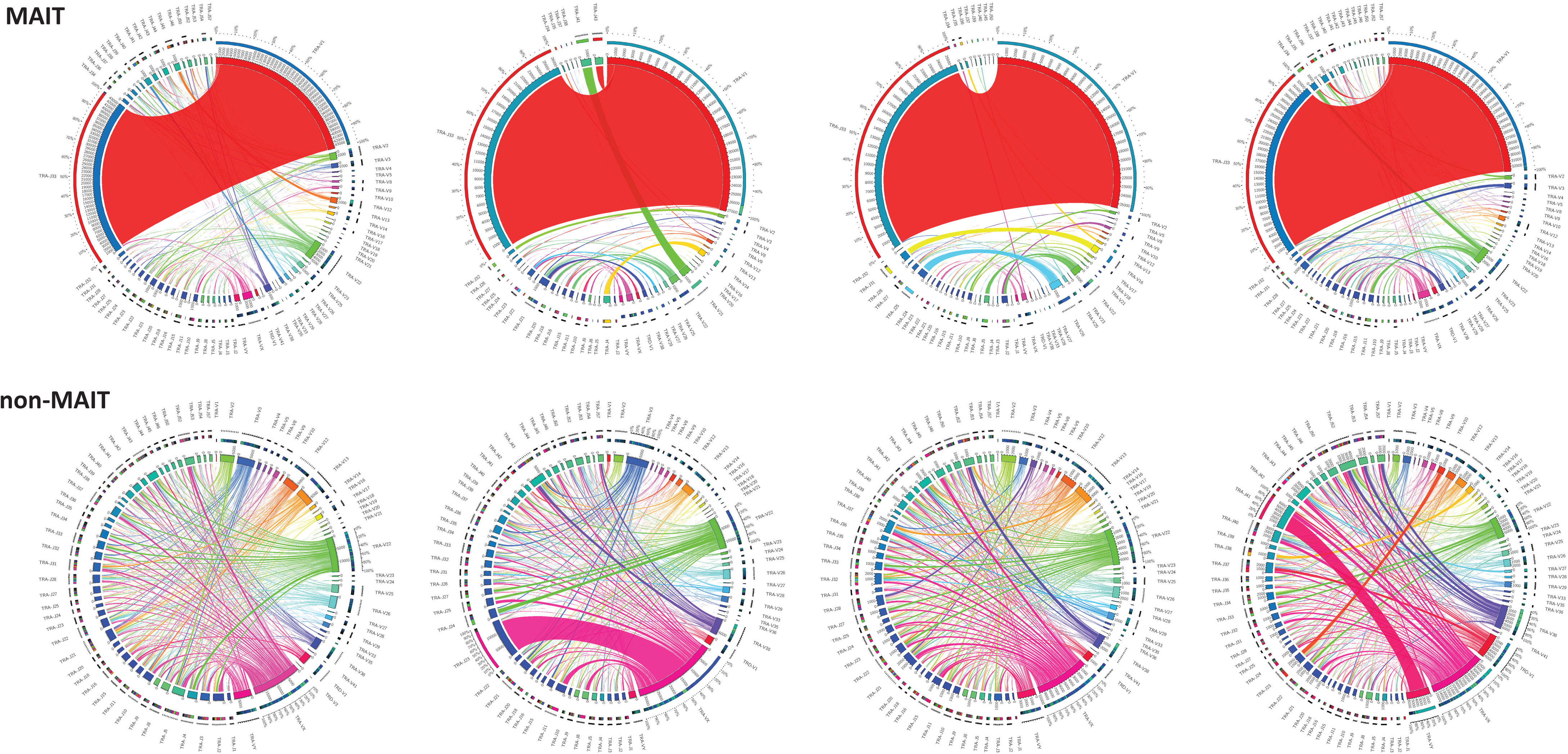
Circos plots showing TRAV-TRAJ combinatorial diversity within MAIT (CD8^+^ human MR1-5-OP-RU tetramer^+^) and non-MAIT (CD8^+^ human MR1-5-OP-RU tetramer^-^) T cells. The inner circle shows the number of sequence reads per V and J gene. TRAV/J combinations are indicated by proportional bands linking the genes using the colour of the V gene. The outer ring shows the frequency of pairing for each TCR gene segment to the reciprocal gene segment coloured according to the paired gene.

**Supplementary figure 4.**
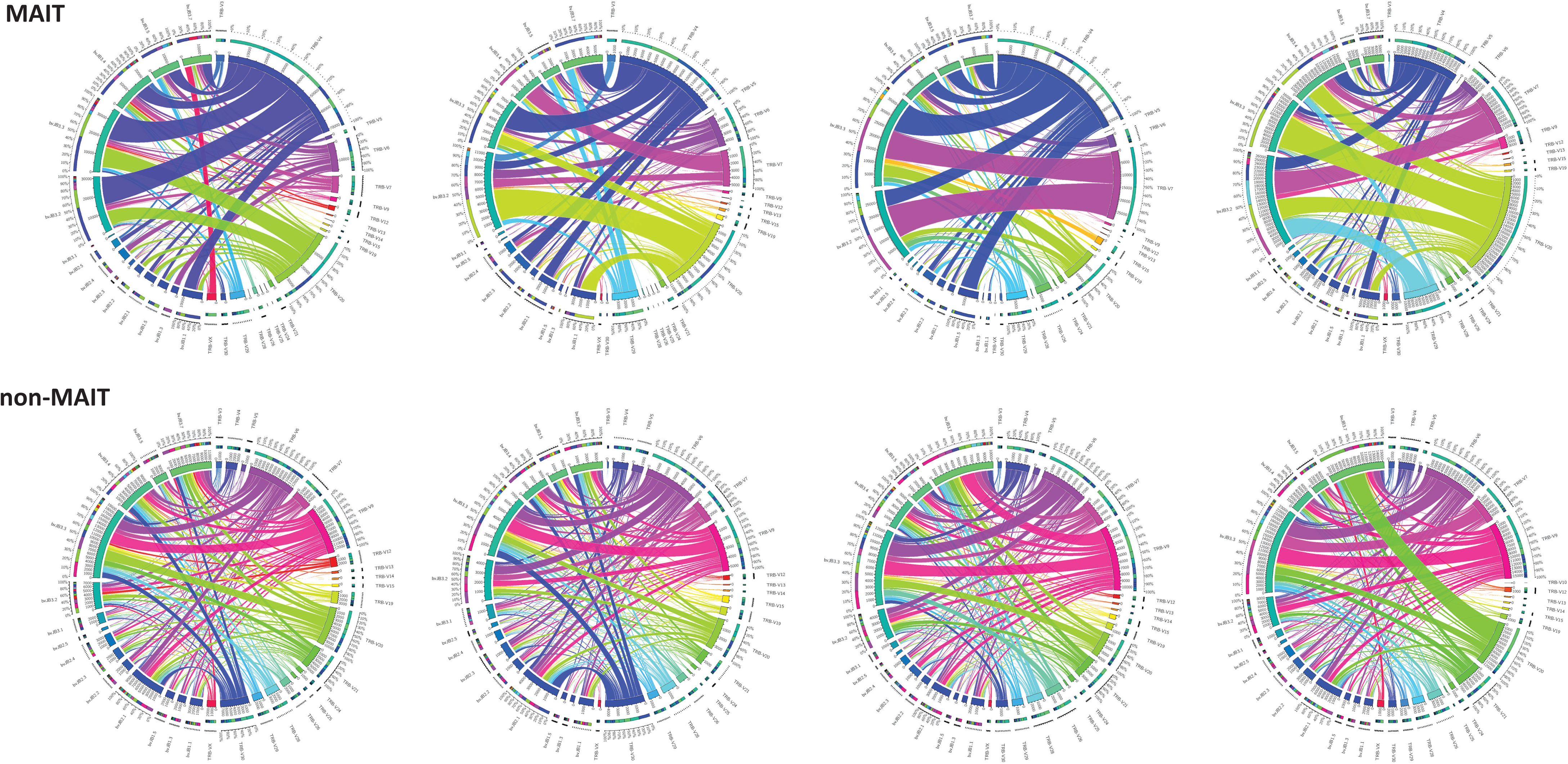
Circos plots showing TRBV-TRBJ combinatorial diversity within MAIT (CD8^+^ human MR1-5-OP-RU tetramer^+^) and non-MAIT (CD8^+^ human MR1-5-OP-RU tetramer^-^) T cells. The inner circle shows the number of sequence reads per V and J gene. TRBV/J combinations are indicated by proportional bands linking the genes using the colour of the V gene. The outer ring shows the frequency of pairing for each TCR gene segment to the reciprocal gene segment coloured according to the paired gene.

**Supplementary figure 5.**
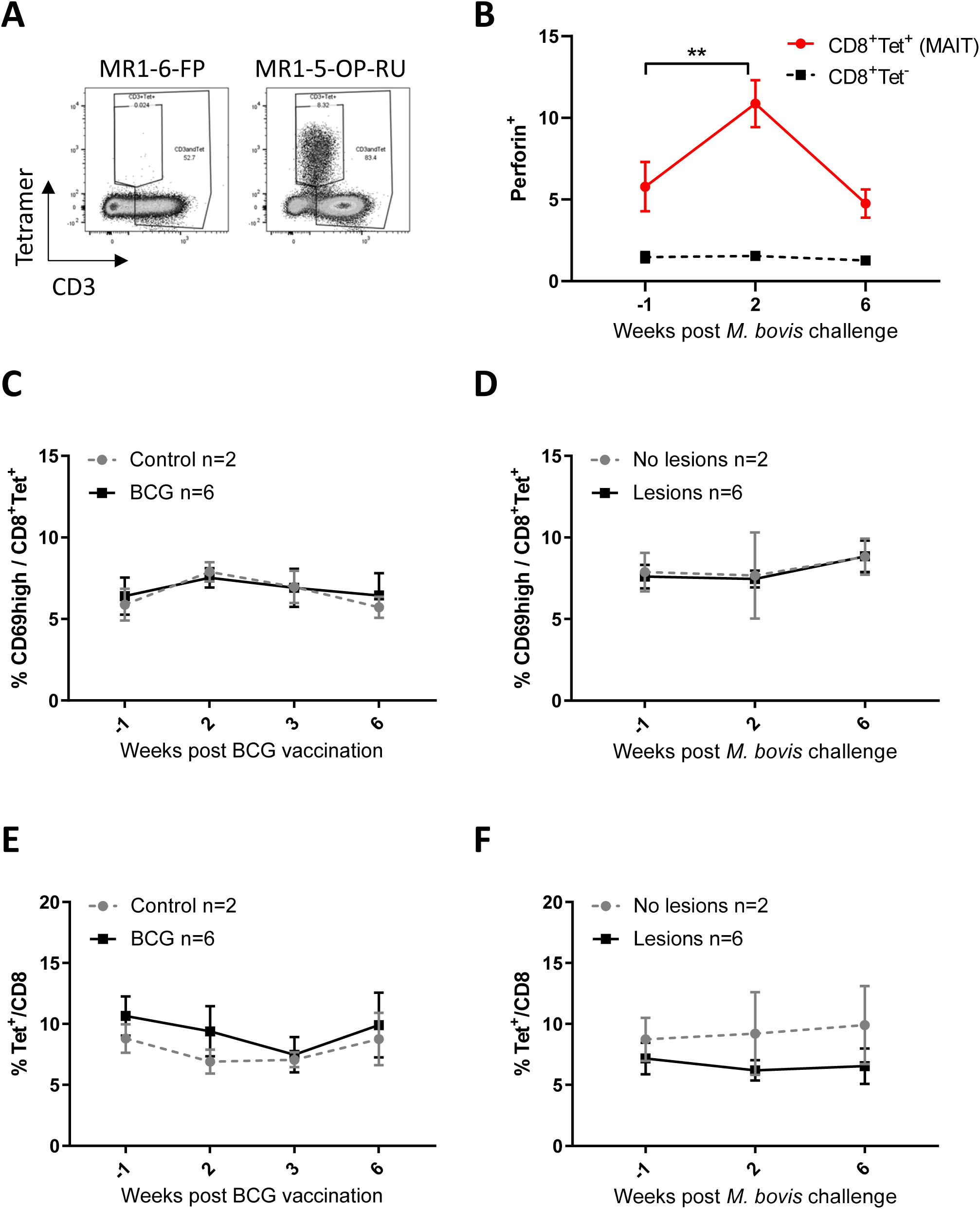
Bovine MAIT cells respond to infections *in vivo*. (**A**) Representative flow cytometry plots of MAIT cell staining in milk. Milk cells were stained with MR1-6-FP or MR1-5-OP-RU tetramer and were gated on FSC&SSC, singlets and live cells. (**B-F**) Characterization of MAIT cells in cattle PBMC following BCG vaccination (**C, E**) and *M. bovis* challenge (**B, D, F**) of calves as in figure 6. Comparison of perforin expression over time by MAIT cells (CD8^+^ MR1-5-OP-RU tetramer^+^) and CD8 T cells (CD8^+^ MR1-5-OP-RU tetramer^-^) within animals with tuberculosis associated lesions in lungs and lymph nodes at post-mortem examination 11 weeks post *M. bovis* challenge (n = 6). (**C, D**) Percentage of MAIT cells (CD8^+^ tetramer^+^) with high CD69 expression. (**E, F**) MAIT cell frequencies within CD8 positive cells in peripheral blood over time. Data were analyzed using two-way ANOVA with time as repeated measures, followed by Sidak’s multiple comparisons post-hoc test comparing changes in perforin expression over time within MAIT cells and CD8 T cells for B and comparing differences between groups at each time point for C-F. Mean ± SEM is indicated in all graphs.

